# Environmental DNA sampling in a terrestrial environment: methods to detect a critically endangered frog and a global pathogen

**DOI:** 10.1101/2020.03.01.968693

**Authors:** Thomas J. Burns, Nick Clemann, Anthony R. van Rooyen, Ben C. Scheele, Andrew R. Weeks, Don A. Driscoll

## Abstract

Environmental DNA techniques have become established as a useful tool for biological monitoring and are used extensively to determine species presence in aquatic systems. However, their application in terrestrial systems has been more limited, likely in part due to difficulties in choosing where to sample and ensuring that collected DNA reflects current species presence. We developed methods to sample eDNA in the terrestrial environment and trialled them under controlled and field conditions. We targeted three species, an elusive critically endangered frog, an abundant non-threatened frog, and the globally distributed amphibian skin pathogen chytrid fungus, which has been implicated in the decline of over 500 amphibian species. We used a sandpaper-sampling surface to ‘trap’ DNA. After sampling, we washed the surface and filtered the wash water to gather material for DNA extraction and subsequent qPCR. Our controlled condition experiments demonstrated that frog and chytrid fungus DNA was detectable after as few as five contacts between a frog and the sampling surface. Furthermore, this DNA remained detectable after two weeks in cool, shaded, outdoor conditions. Our field experiments demonstrated that these techniques were transferable to natural habitats, where we detected both the common and rare amphibian target species, as well as chytrid fungus. Field sampling eDNA results were broadly consistent with those derived from conventional survey methods. Our methods have potential application in non-invasive sampling of amphibians and other species in terrestrial systems, broadening the applicability of eDNA techniques for species detection and monitoring.

## 1. Introduction

Environmental DNA (eDNA) techniques detect organisms from genetic material they shed into the environment- often in the form of skin, mucus, gametes, urine or faeces. They have become established tools for ecologists, particularly for species monitoring where conventional methods may be ineffective, prohibitively labour- or time-intensive, or detrimental to species. They have been used to detect a range of species in mostly aquatic systems, particularly freshwater (Rees et al. 2014; Thomsen & Willerslev 2015; Shaw et al. 2016). For example, they have been used in place of methods that are destructive to habitat (Brozio et al. 2017), to monitor species inhabiting inaccessible areas (Gorički et al., 2017), and, to detect invasive (Ficetola et al., 2008; Valentini et al., 2016) and threatened species (Biggs et al., 2015; Gorički et al., 2017; Olson et al., 2012). They have also been used to detect pathogens (Hashizume et al., 2017; Huver et al., 2015), with a particular focus on amphibian chytrid fungus (*Batrachochytrium dendrobatidis*, henceforth *Bd:* Kirshtein et al. 2007; Walker et al. 2007; Hyman & Collins 2012; Wimsatt et al. 2014). Environmental DNA methods can also be more sensitive or resource efficient than conventional methods (Biggs et al., 2015; Eiler et al., 2018; Lugg et al., 2018; Pilliod et al., 2013; Smart et al., 2015).

Environmental DNA methods have proven an effective tool for monitoring species in aquatic systems. However, many species that are challenging to survey do not occur in aquatic environments or they spend considerable time within terrestrial systems. Use of eDNA methods within terrestrial systems has been more limited and focused mostly on plant and fungal communities (Thomsen and Willerslev, 2015). Recently, application of eDNA methods to detect terrestrial fauna has increased (invertebrates: Bienert et al., 2012; Valentin et al., 2016; Thomsen and Sigsgaard, 2019, vertebrates: Andersen et al., 2012; Kucherenko et al., 2018; Leempoel et al., 2019). However, this has often involved sampling terrestrial fauna from waterbodies (Ishige et al., 2017; Ushio et al., 2017; Williams et al., 2018b) or snow (Franklin et al., 2019b), limiting its spatiotemporal application within the terrestrial environment.

The more common application of eDNA techniques in aquatic systems may be in part due to the relative ease of working with eDNA in aquatic rather than terrestrial systems. Environmental DNA tends to disperse throughout water, and as such, sampling can be less localised. A greater volume of medium can also be included within a single sample from aquatic systems (by passing water over a filter that captures DNA, Rees *et al.*, 2014) compared with terrestrial systems where usually only several grams of soil or sediment can be sampled (Andersen et al., 2012; Bienert et al., 2012). However, dispersal and dilution can complicate the spatial scale of sampling within aquatic systems (Thomsen and Willerslev, 2015). For example, invertebrate DNA was detected > 12 km downstream from its source in a river system (Deiner and Altermatt, 2014).

Within terrestrial systems, there are likely to be fewer opportunities for DNA to be dispersed, increasing the challenge of choosing where to sample, but decreasing the complication of spatial scale. As such, DNA is likely to be concentrated in areas where animals spent time or passed through, potentially requiring a more targeted and precise approach to sampling. Furthermore, DNA degradation rates are generally faster and less variable within aquatic systems (Thomsen and Willerslev, 2015); DNA can become undetectable after days to weeks in water (Dejean et al., 2011; Pilliod et al., 2014), as opposed to potentially years in soils and sediments (Andersen et al., 2012; Barnes and Turner, 2016). As a result, ensuring DNA sampled within a terrestrial environment reflects current species occupancy can be an additional challenge.

Here we detail controlled condition and field experiments of a novel eDNA method for detecting species within the terrestrial environment. We used non-invasive eDNA traps, with a sandpaper sampling surface as focal points to capture DNA from target species that pass over them. Our field system provided ideal conditions for testing our method; variable habitats (wetland vs forest gully) in which to test the robustness of our eDNA trap design and multiple target species, whose presence was confirmed with conventional techniques to evaluate the success of our new terrestrial eDNA sampling technique. Our three target species represent contrasting examples, providing a robust evaluation of our technique: 1) the Baw Baw frog (*Philoria frosti*), is an extremely elusive species with a highly restricted distribution and low adult aquatic association (Hollis, 2011, 2004; Malone, 1985). 2) Amphibian chytrid fungus (*Batrachochytrium dendrobatidis*, henceforth *Bd*), a globally distributed fungus implicated in the decline of hundreds of amphibian species (Scheele et al., 2019). 3) The common eastern froglet (*Crinia signifera*), an abundant generalist (Anstis, 2013) and *Bd* reservoir host (Brannelly et al., 2018; Scheele et al., 2017).

By developing and trialling novel methods for sampling eDNA within terrestrial systems, we expand the potential applications of eDNA techniques as a monitoring tool. More specifically, our method opens up a range of new opportunities to map the terrestrial distribution of *Bd* alongside the movement of amphibians through terrestrial environments where they are otherwise difficult to study. This could enable an understanding of *Bd* distribution, transmission pathways and vectors within the terrestrial environment.

## 2. Materials and methods

### 2.1 Study sites

In our controlled conditions experiment, the abundant frog species *Crinia signifera* were sampled in-situ from a suburban wetland area south-west of Tirhatuan Wetlands, Rowville, Victoria, Australia. Our field experiments using terrestrial eDNA traps were conducted on the Baw Baw Plateau and escarpment area of the Central Highlands of Victoria, Australia.

### 2.2 Controlled conditions experiment

In August of 2016, we established the effectiveness of sandpaper surfaces in trapping and retaining DNA. With gloved hands we caught 23 adult *C. signifera* and prompted them to hop across the surface of a sandpaper sheet. Two grades of sandpaper (80 grit (relatively course) and 400 grit (very fine); *n*=12 and 11, respectively; 28 × 23cm; St Gobain Abrasives, Thomastown, Victoria) and two levels of contact between frog and sandpaper (five and 30 hops; *n*=12 and 11, respectively) were trialled. After sampling, we placed each sandpaper sheet into two zip-lock bags (one inside the other) for storage and transport. We then swabbed each frog following a 35-stroke protocol, with five strokes across each of the centre of the venter, each flank, each thigh and each foot using a fine rayon tipped swab (MWE 100-100, Medical Wire & Equipment, Wiltshire, England). As a field control, we took two sheets of each grade of sandpaper into the field and handled them identically to the treatment sheets, with the exception that we placed no frogs upon them.

We refrigerated 14 sandpaper sheets (12 treatment + two control) at 1-4°C for two days before transfer to a −20°C freezer until DNA extraction. We stored the remaining 13 (11 treatment + two control) at room temperature for 24 hours, then removed them from their bags and placed beneath an outdoor roofed area for two weeks. After this, they were re-packed and stored at −20°C until DNA extraction. The outdoor area was inaccessible to amphibians, sheltered from rain and direct sunlight, but exposed on all remaining sides to the elements. We ensured sandpaper sheets did not contact each other and handled each with a fresh pair of disposable gloves. Weather conditions, measured at the nearest weather station (Ferny Creek, VIC, Australia), were mild and wet during this period, with mean daily rainfall of 4.4mm, and mean daily minimum and maximum temperatures of 8.1 and 13.5°C, respectively (Bureau of Meteorology 2017).

### 2.3 Field experiments

Terrestrial eDNA traps were deployed in two different habitats. First, for 14 days (29^th^ September – 13^th^ October 2016) within montane forest on the Baw Baw Plateau’s escarpment at two *P. frosti* breeding locations (henceforth, gully sites 1 and 2). Second, for 14 days (2^nd^–16^th^ November 2016), at three wetland areas on the Baw Baw Plateau at which *C. signifera* are present and breed (henceforth, wetland sites 1, 2 and 3). Our terrestrial eDNA trap consisted of a sandpaper sheet (as above, 400 grit only) mounted on a black corflute base-plate (290 × 250 × 2.5mm) and pegged flush to the ground, over which a peaked white corflute roof (900 × 400 × 3 and 900 × 600 × 3mm; gully and wetland sites, respectively) was fixed using stakes (Figure 1). We used slightly broader roofs at wetland sites, with the aim of providing additional protection from the elements in this more open environment. Weather conditions during each period, as measured at the nearest weather station (Mt Baw Baw, VIC, Australia), were cool and wet; mean daily rainfalls of 5.5 and 6.4mm, minimum temperatures of −1.3 and 1°C, and, maximum temperatures of 6 and 10°C, respectively (Bureau of Meteorology 2017).

**Figure 1.**
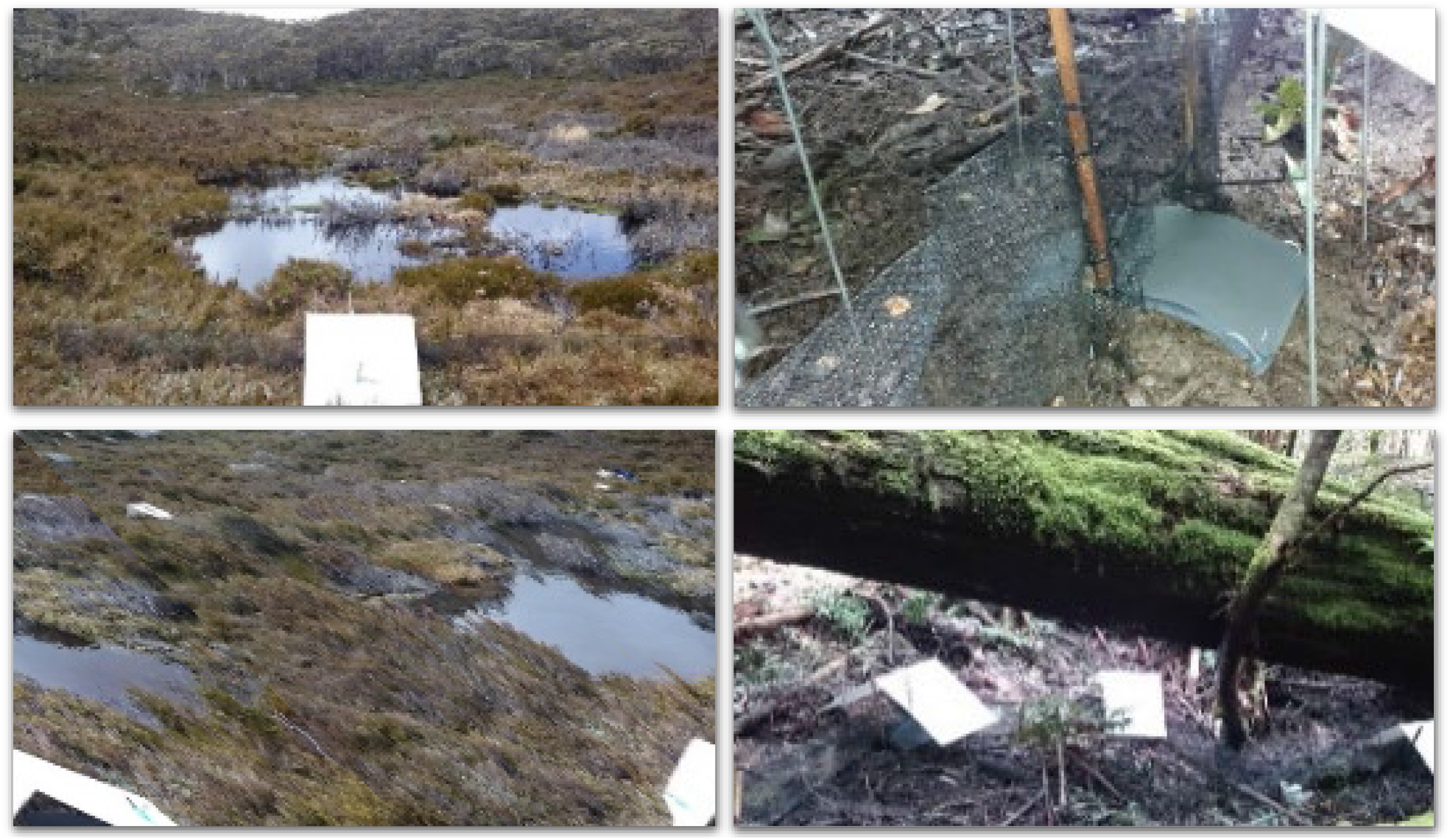
Environmental DNA traps deployed at wetland site 1 (left upper and lower) and at gully site 1 (right upper and lower) in the Baw Baw Plateau and escarpment area, Victorian Highlands, Australia in October and November 2016.

We visited each site multiple times and heard both species calling in their respective habitats, which allowed a qualitative estimation of species presence and abundance. At gully sites, we deployed eDNA traps prior to the peak calling of *P. frosti*, when frogs are expected to be moving into breeding habitat (Hollis, 2011, 2004). At wetland sites, we deployed eDNA traps during the peak activity period of *C. signifera*.

To increase the chance of animal movement over the sandpaper, we deployed drift fencing alongside some traps (henceforth fenced or unfenced traps). Fenced traps at gully sites 1 and 2 (*n*=17 and 7, respectively), were placed over closed pitfall buckets (pitfalls established as part of an unrelated conservation project). Drift fencing enclosed the breeding area from most approaches at site 1, while site 2 was less fully enclosed. Unfenced traps at gully sites 1 and 2 (*n*=17 and 8, respectively) were deployed outside of the drift fenced enclosed area (distance from central ring, mean=6.72m; range=2.7–13.2m), forming an irregular outer ring.

Fenced traps at wetland sites 1, 2 and 3 (*n*=6, 5 and 7 respectively), were deployed with two metres of drift fencing to either side, and placed alternately, but several metres distant from unfenced traps (*n*=6, 6 and 7, respectively). Traps were located either centre, mid-distance from or at the edge of the main *C. signifera* calling areas within wetlands (Figure 1), as determined qualitatively on the day of deployment. We followed strict hygiene sample handling and storage protocols to avoid cross contamination or sample degradation (see appendix S1 for details).

We directly assessed *Batrachochytrium dendrobatidis* (*Bd*) prevalence and mean infection intensity in *C. signifera* at wetland sites for the same period. Sampling was carried out from the 3^rd^-6^th^ of November 2016, by capturing and swabbing *C. signifera* at each site (*n*=32, 18 and 30, at sites 1, 2 and 3, respectively). Due to a lower sample size at site 2 on the first sampling attempt, a second day of sampling was conducted on the 19^th^ of November 2016 (*n*=19) and the combined results used in all analyses. Swabbing followed the protocol described in section 2.2.

### 2.4 Lab processing of samples

We conducted TaqMan^®^ qPCR assays using a Roche LightCycler 480 system (Roche Diagnostics Australia, Castle Hill, NSW, Australia) in a 384-well format. Development of probes and assays for use in qPCR are described in Appendix S2. Reaction volumes were 10μL, containing 5μL of KAPA probe force (Merck), 0.5μL TaqMan^®^ qPCR assay (final primer and reporter probe concentration of 900nmol/L and 250nmol/L, respectively), 2.5μL ddH_2_O, and 2μL of DNA. Each reaction was prepared in triplicate and included two positive controls; a TaqMan^®^ exogenous internal positive control probe (VIC labelled) to test for the presence of inhibition, and DNA extracted from tissue of the target species. Quantitative PCR amplification conditions were 3min at 98°C, followed by 50 cycles of 10s at 95°C and 20s at 60°C. We used the amplification profiles of each PCR to determine the crossing point (Cp) value using the ‘Absolute Quantification’ module of the LightCycler 480 software package. We determined assay efficiency by quantifying DNA extracted from tissue using a Qubit fluorometer (Invitrogen Australia, Mount Waverley, VIC, Australia) and running an assay on a DNA dilution series. The efficiency of all qPCRs was always >95%. We undertook all extractions and qPCR analyses in a room dedicated to low-quantity DNA sources, with qPCR setup undertaken in a laminar flow hood. We added positive controls and standards immediately prior to placing in the Roche LightCycler 480 (separate room). We also included negative controls at all stages (DNA extraction, qPCR) so that we could identify laboratory contamination if present. We found no evidence of contamination in any of the assays.

### 2.5 Interpretation of qPCR results

In the controlled conditions experiment, we ran only a single round of qPCR in triplicate. We designated samples positive if we detected target DNA in two or more wells, equivocal if in one well only, and negative if in none. For the field experiment, we assigned a positive result if the sample returned three positive wells in the first qPCR. Samples that returned one or two positive wells were re-run in a second qPCR, if they returned 2/6 positive wells or more they were designated positive, those with 1/6 positive wells were labelled equivocal. We designated samples that returned no positive wells on the first reaction negative. If we detected inhibition, we applied dilutions until inhibition was no longer detected and assigned the results as above (Figure 2). Swabs of frogs for *Bd* were assigned a positive result only if all three wells were positive.

**Figure 2.**
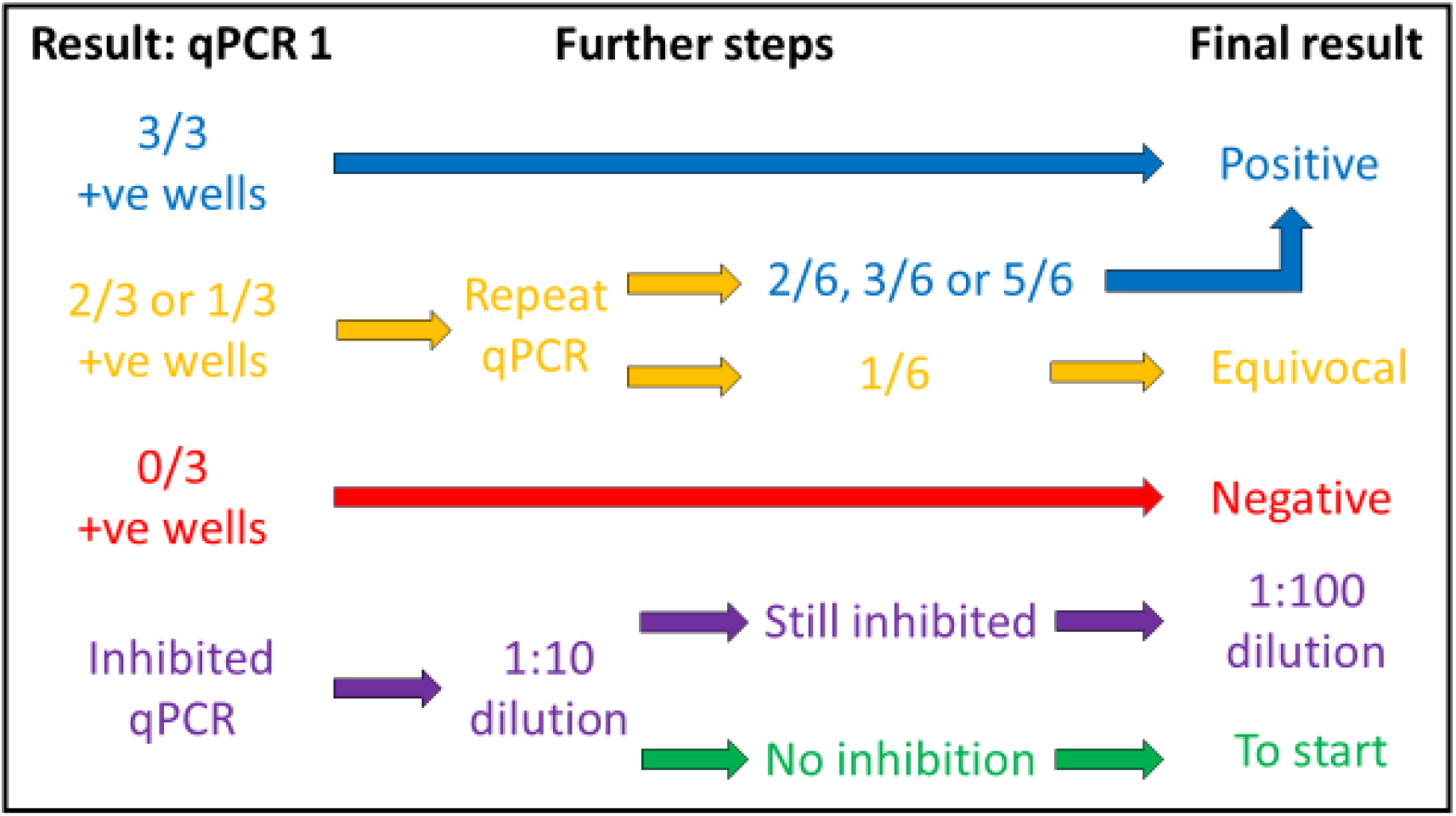
Process for designation of results from qPCR of terrestrial eDNA trap samples in the field experiment.

### 2.6 Statistical analyses

All analyses were carried out using R 3.4.3 (R Core Team 2017). We used ANOVAs (stats package, R Core Team 2018) to examine variation in amount of *C. signifera* DNA detected on sandpaper across each experimental treatment in the controlled conditions experiment. Time of processing, sandpaper grade, and contact with frog were included as categorical explanatory variables. Small sample size (*n*=23) limited the maximum number of model parameters so we modelled each variable individually. The proportion of frogs in the controlled conditions experiment that were *Bd* positive and therefore the number of sandpaper sheets contacted by such frogs was insufficient for analyses of factors affecting *Bd* detection (sandpaper grade, time of processing, contact with frog).

We used binomial generalized linear models with a logit link function to examine factors affecting the detection of *C. signifera* and *Bd* DNA in field experiments at wetland sites. A set of candidate models including all combination of explanatory variables (detailed below), excluding interactions, were produced for each target species and compared using second order Akaike Information Criterion values (AICc) and Akaike Weights (MuMIn package, Barton 2019). We examined parameter estimate confidence intervals from highly ranked models (ΔAICc<2) to determine the influence of fixed effects (MuMIn package).We examined simulated residual plots (DHARMa package, Hartig 2019) to ensure assumptions about the distribution of errors were reasonable.

For *C. signifera* detection: site, trap type (with or without fence) and position in wetland (centre, mid-distance or edge) were included as categorical explanatory variables. To examine *Bd* detection: trap type, position in wetland and *C. signifera* detection on the trap were included as categorical explanatory variables. There were insufficient DNA-positive traps at gully sites for statistical analyses. See Table S2 for a full list of all candidate models.

## 3. Results

### 3.1 Controlled conditions experiment

All skin swab samples collected from *C. signifera* returned positive results for the species’ DNA (*n*=23; mean=275±278.6 SD pg; range=32.8–1,153.3 pg). Eighty-three percent of sandpaper samples contacted by frogs returned at least two positive wells for *C. signifera* (*n*=19; mean=33.5±37.8 pg; range=3.3–176 pg), the remainder returned equivocal results (Table S1). There was a positive effect of frog contact (30 versus 5 hops on surface) on *C. signifera* DNA detected on sandpaper sheets (*F*_1,21_=17.52, *p*=0.0004; Figure 3), but no effect of sandpaper grade or time of processing (*F*_1,21_<0.28, *p*>0.6). Based on direct swabbing, 57% of sampled *C. signifera* were positive for *B. dendrobatidis* (*Bd*) (*n*=13; mean=139±213 SD zoospore equivalent (ZSE); range=3–572 ZSE). Thirty-one percent of sandpaper sheets contacted by *Bd* positive frogs (*n*=13 total) were positive for *Bd* (mean=3±2 SD ZSE; range=1-5 ZSE), 23% equivocal and 46% negative for *Bd* (Table S1). All control sandpaper sheets were negative for *C. signifera* and *Bd*.

**Figure 3.**
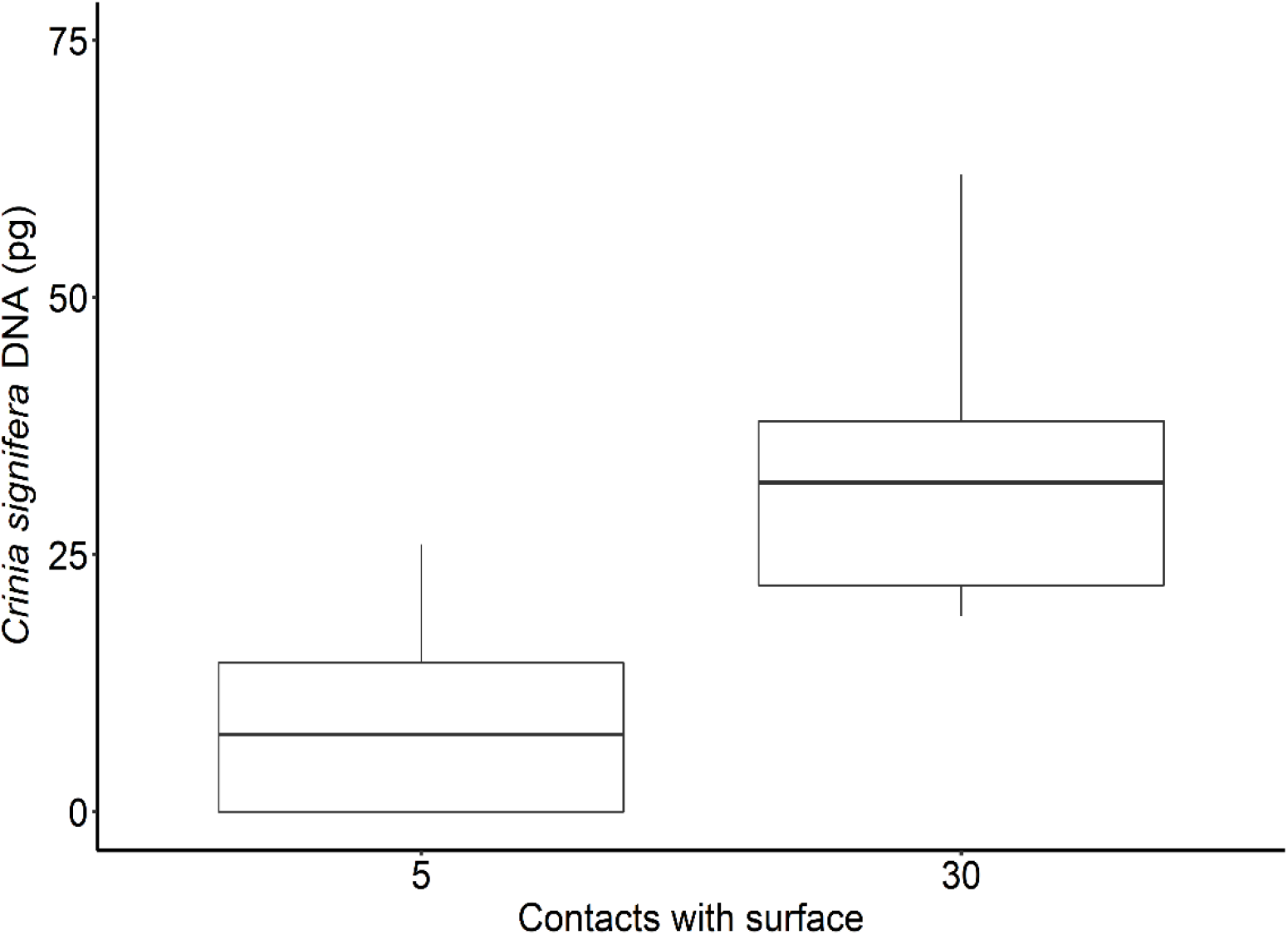
Frog DNA retrieved from eDNA trap sampling surfaces contacted by the frog species *Crinia signifera* during the controlled conditions experiment. Two outliers were removed for clarity, for graph including outliers see Figure S1.

### 3.2 Field experiment

Two of 49 gully site traps (4%) were positive, one trap returned an equivocal result and the remainder were negative for *P. frosti* DNA. No traps in gully sites were positive for *Bd* DNA. Positive results were from fenced traps at gully site 1 (mean=4.5 pg; range=3–5.9 pg), and the equivocal result was from an unfenced trap at the same site (Table 1).

**Table 1.** Summary of results of terrestrial eDNA trapping at wetland and montane gully sites on and around the Baw Baw Plateau, Central Highlands of Victoria, Australia. Target species were Batrachochytrium dendrobatidis, Philoria frosti (gully only) and Crinia signifera (wetlands only). Positive and equivocal (*) results shown for each experimental group with sample size in parentheses. P. frosti only identified at gully sites.

| Habitat | Site | Fenced |  | Unfenced |  |  |  |
| --- | --- | --- | --- | --- | --- | --- | --- |
| **Gully**<br>*(P. frosti)* | **1** | 2 (17) |  | 1* (17) |  |  |  |
|  | **2** | 0 (7) |  | 0 (8) |  |  |  |
|  |  | <u>Centre</u> |  | <u>Mid-distance</u> |  | <u>Edge</u> |  |
|  |  | Fenced | Unfenced | Fenced | Unfenced | Fenced | Unfenced |
| **Wetland**<br>*(Bd)* | **1** | 1 (2) | 1 (2) | 1 (2) | 0 (2) | 0 (2) | 1 (2) |
|  | **2** | 0 (2) | 0 (3) | - | - | 0 (4) | 0 (4) |
|  | **3** | 2 (3) | 0 (2) | 0 (2) | 0 (3) | 0 (2) | 1 (2) |
| **Wetland**<br>*(C. signifera)* | **1** | 2 (2) | 2 (2) | 1 (2) | 1 (2) | 1 (2) | 1 (2) |
|  | **2** | 1 (2) | 1 (3) | - | - | 2 (3) | 1 (3) |
|  | **3** | 3 (3) | 2 (2) | 1 (2) | 0 (3) | 0 (2) | 0 (2) |

Fifty-one percent of the traps placed within wetlands (*n*=19/37) were positive for *C. signifera* DNA (mean=17.2±43.9 SD pg; range=1.76–191.8 pg), and 19 % were positive for *Bd* DNA (mean=2.9±0.8 ZSE; range=1.8–4.1 ZSE, Table 1). *Crinia signifera* positive traps were obtained from all sites (67, 45 and 43 % positive traps at sites 1, 2 and 3, respectively), from both fenced and unfenced traps, located in all areas of the wetlands sampled (Table 1). Two models had ΔAICc<2, both included position in wetland and one also included trap type (Table 2). Parameter estimates indicate trap position was important for *C. signifera* detection, as 95% confidence intervals exclude zero (mid-distance, −4.148 to −0.199; and edge, −3.88 to −0.335, Table S2), with greater detections at traps located in central positions. Trap type was not a useful predictor of *C. signifera* detection (−2.465 to 0.542, Table S2).

**Table 2.**
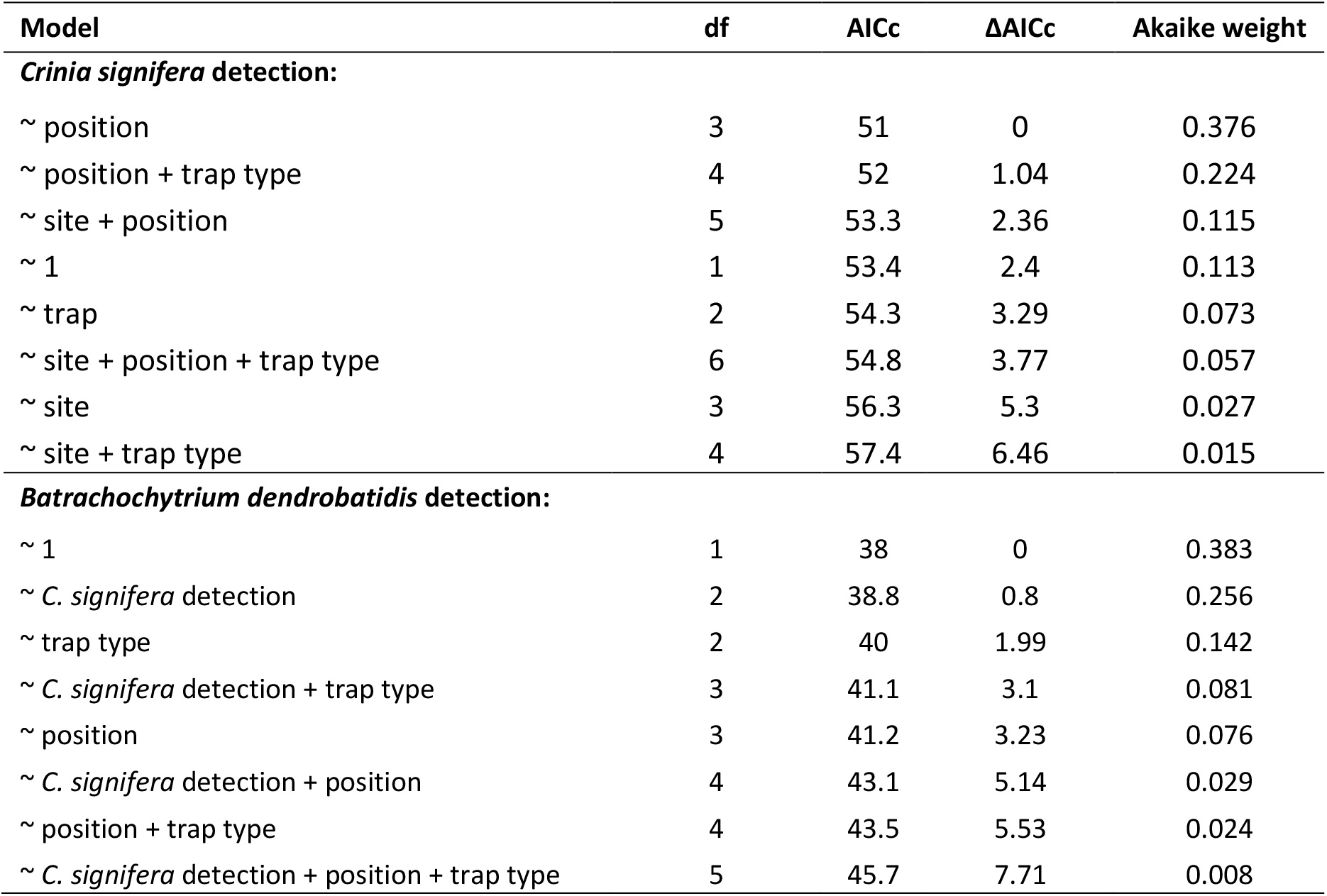
Candidate models used to examine factors influencing detection of *Crinia signifera* and *Batrachochytrium dendrobatidis* (Bd) on terrestrial environmental DNA traps deployed in the Baw Baw plateau area of south-eastern Australia in 2016. Models are ranked by second order Akaike Information Criterion (AICc) and delta AIC (ΔAIC)

*Batrachochytrium dendrobatidis* positive traps were from wetland sites 1 and 3 only (25 and 21% traps positive, respectively), and were both fenced and unfenced (Table 1). The majority of positive traps (57%) were located in central areas of wetlands (central, *n*=4; mid-distance, *n*=1; edge, *n*=2). Five of seven *Bd* positive traps were also positive for *C. signifera*. Three models, including the null model, had ΔAICc<2. High-ranking models included *C. signifera* detection and trap type as explanatory variables (Table 2). Parameter estimate 95% confidence intervals indicate neither were good predictors of *Bd* detection: *C. signifera* detection (−0.739 to 2.839) and trap type (−2.081 to 1.239, Table S2). Estimated *Bd* detection in our field experiment, calculated using population *Bd* prevalence and *C. signifera* detection on traps, was 42% (see Appendix S3 for calculation) and comparable with *Bd* detection in our controlled condition experiment (31%).

*Batrachochytrium dendrobatidis* prevalence of swabbed *C. signifera* was high across sites 1, 2 and 3 (*n*=32, 37 and 30; prevalence = 91, 80 and 93%, respectively). However, infection intensity of swabbed frogs was relatively low (mean=76.3±106.5; 134.2±245.1 and 183.9±204 SD ZSE, at sites 1, 2 and 3, respectively) compared to published values for *C. signifera* in upland areas of south-eastern Australia (Brannelly et al., 2018).

## 4. Discussion

Innovative approaches are required to apply eDNA techniques to monitor fauna in terrestrial systems. We used a sandpaper surface to sample contemporary eDNA from amphibians that have moved over a defined spatial area within the terrestrial environment. Detection rates of *C. signifera,* a common frog species, were high with 83% positive (remainder equivocal) in our controlled condition experiment, and we detected the amphibian skin pathogen *Batrachochytrium dendrobatidis* (*Bd*) with a success rate of 30-42%. Our results were similar to eDNA studies that targeted individual animals, whose presence was confirmed using other methods, from snow and soil in wild settings (Franklin et al., 2019, 81% samples ≥2 positive wells; Kucherenko et al., 2018, 67%). However, our detection success was lower than values reported for species presence in captivity, where almost all species present were detected (Andersen et al., 2012; Ushio et al., 2017). Our technique provides a step towards systematic sampling of the terrestrial landscape for eDNA, in a manner akin to how audio recorders, cameras and pitfall traps are used to sample sounds, images and animals. To our knowledge, this is the first use of eDNA traps to detect both a terrestrial species and a skin pathogen they carry.

Our controlled experiment results suggest high detection of frogs that interacted with traps; individuals were detectable from as few as five contacts, two weeks later (Table S1). Furthermore, the amount of DNA detected was greater when there were more contacts (Figure 3), suggesting that there may be potential for estimating relative abundance by ranking traps based upon amount of DNA.

We were unable to monitor frog interactions with our field experiment traps to calculate exact detection rates; however, results from our field experiment were similar to those achieved using aural survey methods. At wetland sites, detection of *C. signifera* was generally high (51% of all traps), but greater at centrally located traps (79%), where the highest density of calling frogs were present. At our encircled gully sites, aural surveys of *P. frosti* were carried at the beginning and end of our eDNA trapping period as part of an unrelated study. At site 1, one and then four males were recorded respectively, and at site 2, one male was recorded calling each time (D. Gilbert, personal comment). This suggests a low level of frog movement across our traps and relatively high frog DNA detection rates (two positive traps at site 1).

Our method was less sensitive at detecting *Bd* DNA left by *Bd* positive frogs; 31% and an estimated 42% detection success in controlled condition and field experiments, respectively (Appendix S3). However, we note the infection loads of *Bd* positive *C. signifera* in both experiments, determined by extensive swabbing of the ventral surface of the body, were also relatively low compared with previously published values for this species (Brannelly et al., 2018). Given the high *Bd* prevalence within *C. signifera* populations and the results of our controlled condition experiments, we expected coincident detection of *Bd* and frog DNA in our field experiments- yet two of seven *Bd* positive traps were negative for *C. signifera*. It is unlikely that other amphibian species were responsible for this result; across four visits to each of these sites between November 2016 and April 2017, we detected only *C. signifera* during extensive daytime searching of ground vegetation for amphibians (T. Burns, personal observation). Assuming similar retention, degradation and detection rates for *Bd* and *C. signifera* DNA, this may indicate that zoospores reached the sampling surface without carriage upon an amphibian. Potential alternatives include zoospores being dispersed by the elements (Kolby et al., 2015a), or, non-amphibian vectors or hosts (Garmyn et al. 2012; Kilburn et al. 2011; McMahon et al. 2013; Brannelly et al. 2015; Shapard et al. 2011). We detected no *Bd* from gully site traps, a result consistent with *Bd* swabbing carried out at these sites in another study that immediately followed our own (*n*=16 *P. frosti*, all *Bd* negative; D. Gilbert, personal communication).

Our methods have potential application in the systematic sampling of species within the terrestrial environment. Here, they offer some potential advantages over both previously applied eDNA methods targeting terrestrial fauna and conventional survey techniques. Currently, eDNA methods targeting terrestrial fauna tend to sample substrate or focal water sources (Andersen et al., 2012; Ishige et al., 2017; Leempoel et al., 2019; Williams et al., 2018). By deploying a sampling surface, our methods are less susceptible to potential issues with the temporal scale of sampling associated with more variable and potentially slower DNA degradation rates in terrestrial substrates (Thomsen and Willerslev, 2015). Furthermore, they are not spatially limited by the availability of water sources or to the terrestrial species that can be detected from them (Ushio et al., 2017). In comparison with conventional techniques, our methods are less explicitly restricted to particular life history stages, periods in the year or times of day, as for example aural surveys for amphibians.

As well as the broad applications outlined above, we also wish to highlight a specific application for which our methods may be particularly suited. Amphibians with low aquatic associations account for 28% of the 500 species impacted by *Bd*, with most of these species experiencing severe and ongoing declines (Scheele et al., 2019). Yet, studies examining *Bd* in the wild have been for the most part restricted to sampling water bodies or amphibians directly (see Kolby et al., 2015b for terrestrial *Bd* exception). Similarly, studies examining amphibian habitat use have tended to focus on more accessible, often aquatic, breeding congregations. As a result, understanding of the distribution and dispersal of *Bd*, and of amphibian movements within the terrestrial environment is often lacking.

Moderate detection rates of frog DNA on traps at the drier edges of wetland sites (43%) suggest our technique may be valuable for sampling terrestrial areas where animals are present at lower densities. We suggest that our methods may offer a potential sampling approach for determining *Bd* distribution within the terrestrial environment. Furthermore, by examining species associated with *Bd* detections, it may be possible to identify potential vectors or reservoirs of disease.

A logical next step in further developing our terrestrial eDNA trapping methods would be to use them in conjunction with camera traps. This would allow quantification of animal-trap interactions for direct calculation of detection rates in wild settings and assessment of species that can be targeted using this method. Furthermore, it would allow targeted sampling of individuals of interest. For example, to confirm species identification of animals difficult to identify from images, as demonstrated with eDNA collected from snow (Franklin et al., 2019b). It could also be used to extract additional information about a population or area, such as disease presence, as has been done using eDNA techniques in aquatic systems (Hashizume et al., 2017; Huver et al., 2015), including with *Bd* (Wimsatt et al., 2014, among others). Finally, there are other settings were our methods have potential application, for example to improve detection or reduce the number of site visits required when surveying reptile tile grids.

At gully sites, water splashes from the muddy substrate led to the soiling of sampling surfaces, and most samples required dilution to overcome qPCR reagent inhibition. This could have two main impacts on our results. First, dilution decreases the probability of detecting DNA present in samples at low concentrations (McKee et al., 2015), and second, soiling increases the possibility that *P. frosti* DNA shed into the environment at an earlier date could be transferred onto the sampling surface. However, we think it unlikely that the temporal scale of sampling was altered, as DNA shed by amphibians onto the soil surface in this environment is likely to degrade rapidly (Kucherenko et al., 2018; Walker et al., 2017). We suggest modification of our trap design to include a broader, more enclosing roof structure or attaching a fringe of shade cloth to the trap base mount to reduce muddy splashes and subsequent inhibition.

## Conclusion

We demonstrate that small-bodied amphibians and a globally important fungal pathogen are detectable in the terrestrial environment using eDNA methods. Our methods have application for monitoring species in terrestrial environments or supplementing other survey methods to provide additional information. In particular, our methods could be useful for species difficult to detect using conventional methods. Furthermore, our methods have specific application for examining *Bd* distribution, transmission pathways and vectors within the terrestrial environment.

## Supporting information

Electronic supplementary material

## 5. Acknowledgments

We thank Parks Victoria and the Victorian Government’s Department of Environment, Land, Water and Planning for permits to carry out fieldwork, and the Mount Baw Baw Alpine Resort for providing accommodation throughout fieldwork. We are grateful to David De Angelis, Deon Gilbert, Jet Black and Timothee Poupart for assistance in the field. We also thank technical staff at Deakin University-Clorinda Schofield, Tom Schneider and Tom Healey for assistance with trap design and field preparation. Furthermore, we thank our funders, the Holsworth Wildlife Research Endowment and the Centre for Integrative Ecology, Deakin University, without which this work would not have been possible.

Research was approved by the Deakin University Animal Ethics Committee (projects B29-2015 and B27-2016) and conducted under permits (# 10007793 and 10008072) issued by the Victorian Government’s Department of Environment, Land, Water and Planning.

## 6. Author contributions

TJB, DAD, NC, ARVR, BCS and ARW conceived the ideas and designed methodology; TJB conducted fieldwork; TJB and ARVR conducted lab-work; TJB analysed the data and led the writing of the manuscript. All authors contributed critically to the drafts and gave final approval for publication.

