## Supplementary material for "Environmental DNA sampling in a terrestrial environment: methods to detect a critically endangered frog and a global pathogen": Electronic supplementary material

**Electronic supplementary material (ESM)**

**List of included supplementary materials**

**Appendix S1.** Description of hygiene and sample handling and storage protocols followed.

**Appendix S2.** Development of probes and assays for qPCR assays.

**Appendix S3.** Estimation of *Batrachochytrium dendrobatidis* detection in field experiments.

**Table S1.** Controlled conditions experiment results summary.

**Table S2.** Parameter estimates for factors explored to be influencing the detection of *Crinia signifera* and *Batrachochytrium dendrobatidis* (*Bd*) on terrestrial environmental DNA traps.

**Figure S1.** Frog DNA retrieved from eDNA trap sampling surfaces contacted by the frog species Crinia signifera during the controlled conditions experiment. *represents outliers.

**Appendix S1.** Description of hygiene and sample handling and storage protocols followed.

Stakes, corflute mounts and roofs were disinfected with a 1% sodium hypochlorite solution prior to use and were each deployed at only one site. Equipment used at multiple sites was thoroughly cleaned, soaked in a 1 % sodium hypochlorite solution for a minimum of one hour, rinsed with fresh water and allowed to dry before re-use. A fresh pair of nitrile gloves was worn when handling each sandpaper sheet and the staple gun used to attach them to the corflute mount. The staple gun was handled with care, ensuring it did not contact any surface other than the sandpaper sheet. Between uses it was stored within a sealed zip-lock bag. As a field control for cross contamination a sheet of sandpaper was taken to each field site during deployment and retrieval of traps. These were handled identically to those in treatment groups, but were not attached to traps; during the two week deployment period control sheets were stored at 1 - 4 °C. After retrieval, sandpaper sheets were placed into two zip-lock bags (one inside the other) and refrigerated at 1 - 4 °C for four days, before being transported in a cooler from our field site and transferred to a -20 °C freezer.

**Appendix S2.** Development of probes and assays for qPCR assays

We PCR amplified and sequenced a fragment of the mitochondrial NADH gene from tissue samples of *C. signifera* and *P. frosti* from the study area. We then compared these sequences with those available from each species (and other related species) on GenBank (NCBI; www.ncbi.nlm.nih.gov) to design species-specific quantitative PCR (qPCR) assays. For sequencing, tissue samples (toe clips) were collected from ten *C. signifera* from the Baw Baw Plateau area and swabs were obtained from two *P. frosti* held at Zoos Victoria (Parkville, Victoria, Australia) that had been collected from the same area. We extracted DNA from toe clip samples using a QIAGEN DNeasy kit (Qiagen Pty Ltd, Victoria, Australia). Swabs were extracted with 3µL proteinase K (Qiagen Pty Ltd, Chadstone, Victoria, Australia) and 197µL of 5% Chelex® solution, they were incubated at 55°C for 1hour and 95°C for 15min with periodic vortexing. Samples were stored at -20°C until use in qPCR. Prior to qPCR, samples were centrifuged at 12,000g for 2min and supernatant from just above the Chelex® resin was used for qPCR. PCR amplification was undertaken with primer ND4 TGACTACCAAAAGCTCATGTAGAAGC and Limno3 TRTGTTCNCGGSTTGTWGT for *C. signifera* and ND4 and Limno2 TRTGTTCNCGGSTTGTWGT for *P. frosti* (Arevalo et al., 1994; Schäuble et al., 2000) to amplify 616 bp or 791 bp of the NADH dehydrogenase subunit 4 (ND4) gene for each species.

PCR amplifications were prepared in 20 µL volumes for each sample containing 6.4 µL ddH2O, 0.8 µL each primer pair (10 µM), 10 µL of Qiagen multiplex mastermix (Qiagen), and 2µl DNA extract. PCR cycling conditions were 95 °C for 15 minutes, 35 cycles of 94 °C for 30 seconds, 48 °C for 30 seconds, and 72 °C for 90 seconds, with a final extension step of 72 °C for 10 minutes on an Eppendorf Gradient *S* Master Cycler (Eppendorf). PCR products for each sample were directly sequenced on an ABI3730 XL DNA sequencer (Applied Biosystems) in both directions using the primers described above. Sequences were aligned and manually edited in Geneious version 7.1.9 (Biomatters Ltd., Auckland, New Zealand). Unique haplotypes were submitted to Genbank (accession numbers awaiting approval).

Species-specific TaqMan® copy number assays (Thermo Fisher Scientific Inc. Waltham, MA) were developed to target mitochondrial NADH dehydrogenase 4 (ND4) gene fragments of 95 and 60 base-pair (bp) for *C. signifera* and *P. frosti* respectively. *Crinia signifera* (BawBaw) forward *5′*-GCGGTTACGGCATCTTACGA-3′, reverse 5′-GACCACGCCGCATATGG-3′ and probe 5′-CATGAATCTTTGTATCCATTAGCC-3′ and *P. frosti* forward 5′-CGAGGAGAATGTGTTCTCGGG-3′, reverse 5′-ACCTCCCGCCCCACTTAA-3′ and probe 5′-AATCTACCCCCCCCACAATA-3′. We tested the specificity of each assay on the DNA extracted above for each species and on DNA from other amphibians detected or known from the study area (*Litoria verreauxii alpina* and *Litoria ewingii*). We confirmed the specificity of each assay by checking that only the target species was amplified in each assay respectively. For *Bd* the TaqMan® assay and standards were as described in Boyle et al., 2004. For the controlled conditions experiment we used the *C. signifera* qPCR described in Guillera-Arroita et al., 2017; the *C. signifera* marker described above was subsequently developed to target individuals from the Baw Baw plateau due to geographic differences in haplotypes for the ND4 gene.

**Appendix S3.** Estimation of *Batrachochytrium dendrobatidis* (*Bd*) detection in field experiments.

$$Bd\ \text{detection} = \frac{\text{Observed } Bd \text{ positive traps}}{\text{Expected } Bd \text{ positive traps}}$$

$$\text{Expected } Bd \text{ positive traps} = \text{total traps deployed } \times \text{proportion of traps frog positive } \times \text{population } Bd \text{ prevalence}$$

$$Bd\ \text{detection} = \frac{7}{(37 \times 0.51\ \times 0.88)} = 0.42$$

**Table S1.** Sandpaper sheet samples positive for *Crinia signifera* and *Batrachochytrium dendrobatidis* (*Bd*) in controlled condition experiment. Sheets considered positive for when a minimum of 2/3 wells return positive results, only sheets contacted by *Bd* positive *C. signifera* (swabs returning 3/3 Bd positive wells) considered within *Bd* group. Sample size of each group indicated in brackets. Coarse and fine sandpaper were 80 and 400 Grit, respectively.

| Target | Processing | Sandpaper | Positive sheets (n) | |
|---|---|---|---|---|
| | | | 5 contacts | 30 contacts |
| *Crinia signifera* | Immediate | Coarse | 3 (3) | 3 (3) |
| | | Fine | 2 (3) | 3 (3) |
| | After two weeks outside | Coarse | 1 (3) | 3 (3) |
| | | Fine | 2 (3) | 2 (2) |
| *Batrachochytrium dendrobatidis* | Immediate | Coarse | 0 (2) | 0 (2) |
| | | Fine | 1 (1) | 1 (2) |
| | After two weeks outside | Coarse | 0 (1) | 1 (2) |
| | | Fine | 0 (1) | 1 (2) |

**Table S2.** Parameter estimates for factors influencing the detection of *Crinia signifera* and *Batrachochytrium dendrobatidis* (*Bd*) on terrestrial environmental DNA traps. We extracted parameter estimates from highly ranked model (ΔAICc<2) in each candidate model set. Model reference levels, for *C. signifera* detection: position = centre of wetland; trap type = fenced; and site = 3, and for *B. dendrobatidis* detection: *C. signifera* detection = negative; position = centre of wetland; and trap type = fenced. CI, 95% confidence intervals; SE, standard error.

| Parameter; level | Estimate | SE | Lower CI | Upper CI |
|---|---|---|---|---|
| ***C. signifera* detection:** | | | | |
| Position; edge | -1.975 | 0.886 | -3.711 | -0.238 |
| Position; mid-distance | -2.033 | 0.987 | -3.968 | -0.098 |
| Trap type; fenced | -0.902 | 0.753 | -2.378 | 0.574 |
| ***Bd* detection:** | | | | |
| *C. signifera* detection; positive | 1.05 | 0.913 | -0.739 | 2.839 |
| Trap type; fenced | -0.421 | 0.847 | -2.081 | 1.239 |

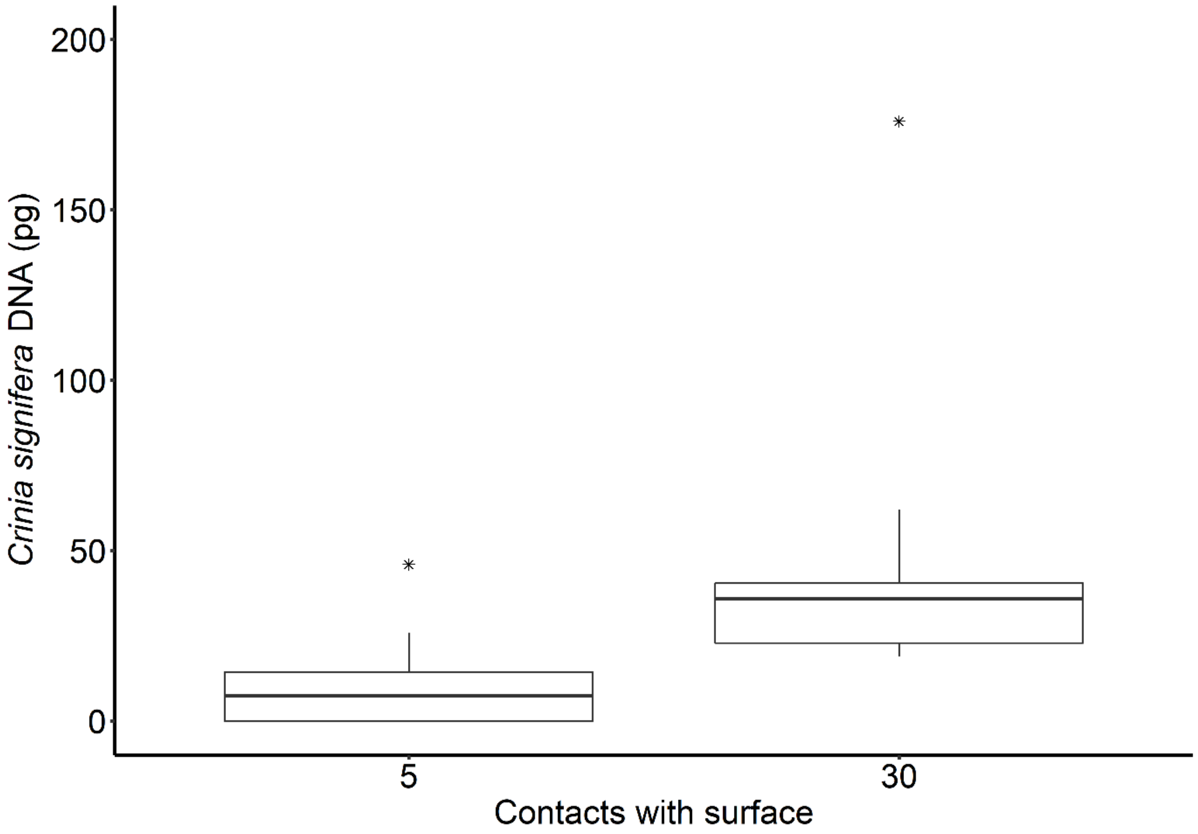


**Figure S1.** Frog DNA retrieved from eDNA trap sampling surfaces contacted by the frog species *Crinia signifera* during the controlled conditions experiment. * represents outliers.
